# Comparative benchmarking of CRISPRi and CasRx in standardized pluripotent stem cell platforms reveals context-dependent knockdown performance

**DOI:** 10.64898/2026.05.13.724469

**Authors:** Lu Ni, Takuya Murakami, Satoshi Suzuki, Mari Hamao, Michiko Nakamura, Chikako Okubo, Kazutoshi Takahashi

## Abstract

Advances in transcriptome profiling have revealed transcriptomic differences across different cellular states. However, functional interpretation requires precise perturbation tools and experimental frameworks. This study benchmarked two widely used modalities: CRISPR interference (CRISPRi) and Cas13d/CasRx. A standardized workflow was established to generate human pluripotent stem cells (PSCs) with inducible ZIM3-dCas9 or CasRx expression. The cell lines were subjected to flow cytometry, copy number, and immunocytochemical analyses. The knockdown performance was validated via robust OCT4 suppression and the expected downstream effects on pluripotency genes. Time-course measurements indicated that CRISPRi produced faster and stronger repression but slower recovery after inducer withdrawal. In contrast, CasRx yielded slower and typically weaker knockdown with rapid reversibility. Furthermore, a key limitation of CRISPRi was demonstrated using the *ATF5*-*NUP62* locus, wherein CRISPRi could co-repress genes with overlapping promoter regions. In contrast, CasRx avoids these limitations and supports isoform-resolved targeting of circular and alternatively spliced transcripts, albeit with variable efficiency. These results provide practical guidance for selecting complementary knockdown tools to improve the interpretability of transcriptomic function studies.

**MOTIVATION:** Advances in transcriptome profiling have enabled the detection of subtle cell type-specific differences. However, mechanistic interpretation still depends on perturbation tools that can modulate transcripts with high precision and efficiency. Recent CRISPR-based modalities, CRISPRi and Cas13/CasRx, function as robust and orthogonal methods to achieve the knockdown of specific gene targets. However, a standardized approach for cell line preparation and comparative studies on their relative performances and limitations remains unclear. Consequently, this study presents a standardized workflow for generating cell lines that support high-efficiency knockdown using CRISPRi and CasRx. Moreover, it compares the trade-offs in potency, reversibility, and isoform resolution, along with a practical overview of method-specific pitfalls to guide tool selection and data interpretation in future studies.

**HIGHLIGHTS:**

- Doxycycline-inducible *AAVS1* knock-in human PSC platforms for CRISPRi (ZIM3-dCas9) and CasRx (RfxCas13d) were generated to enable standardized RNA perturbation experiments.
- The prepared cell lines demonstrated strong OCT4 knockdown, with expected downstream effects on the expression of another pluripotency gene, *NANOG*.
- A comparison of knockdown characteristics and their reversibility revealed rapid and sustained repression with CRISPRi, whereas slow but rapid recovery was observed with CasRx.
- A CRISPRi-specific off-target effect arising from TSS proximity/overlap (*ATF5*-*NUP62*) was identified, whereas CasRx achieved *ATF5* knockdown without collateral repression of the neighboring *NUP62* gene.
- CasRx enables isoform-resolved knockdown of structural isoforms (circHIPK3 vs. linear HIPK3 mRNA) and splice isoforms (RAB6A-iso1 vs. RAB6A-iso2).

## INTRODUCTION

The cellular transcriptome encodes most of the information that dictates the functional and behavioral characteristics of an organism. Significant technological advancements have been observed in our ability to map the cellular transcriptome over the past three decades, ranging from microarrays^1^ to next-generation sequencing and single-cell approaches^2^. These techniques have transformed our ability to track and analyze transcriptomic changes that are implicated in cellular function and disease progression. Despite these advances, tools capable of actively modulating specific transcripts are required to fully elucidate the functional dynamics of individual transcripts and establish causal relationships between transcriptional changes and cellular phenotypes. These tools must provide highly specific and programmable control of gene expression and facilitate the regulation of multiple genes across diverse organisms on a genome-wide scale.

Among the currently available methods, RNA interference (RNAi) is a simple yet effective way to change specific transcript abundance and validate the function of specific transcripts without modifying the host genome^3^. Small interfering RNA (siRNA) is currently regarded as the gold standard,^4^ as it only requires the design and introduction of a short (19-23 nt) dsRNA to initiate the knockdown (KD) of the target transcript. However, in practice, significant off-target effects have been reported due to crosstalk with the microRNA machinery.^5^ Moreover, the effectiveness of RNAi is limited by the accessibility of the target transcript to the silencing machinery, whereby RNAi efficacy is significantly reduced when targeting highly structured transcripts^6^.

CRISPR-Cas is a diverse family of nucleases that rely on the presence of a guide RNA (gRNA) for guidance and targeting specificity^7,8^. The gRNA comprises a structured spacer sequence segment, which facilitates the association with a specific Cas nuclease, and a non-structured spacer sequence segment that recognizes the target nucleic acid sequence^9^. As such, the appropriate design of the spacer sequence and selection of the Cas nuclease type confer CRISPR/Cas systems the flexibility to target various nucleic acids. Although initially used in genomic engineering^10,11^, rapid developments and recent discoveries in CRISPR/Cas technologies have enabled new methodologies for precise targeting and reversible modulation of transcript expression, without perturbing host genomic sequences^12–17^.

CRISPR interference (CRISPRi) has emerged as a highly effective method for the selective silencing of target transcripts with minimal off-target effects^12,16^. The KRAB-dCas9 uses a catalytically dead CRISPR-Cas9 protein (dCas9) fused with the transcriptional repressor Krüppel-associated box (KRAB) domain to reduce the transcription of the target gene by triggering heterochromatin-mediated gene silencing at the transcription start site (TSS) ^3,16^. This grants CRISPRi single-gene level precision and a potent phenotype compared to traditional RNAi, often almost completely repressing the expression of the target gene (>99%)^16,17^. As such, CRISPRi has gained wide application as an alternative method to siRNA silencing.

Recently, the discovery of a new CRISPR-Cas13 family of nucleases has introduced a novel vector into direct transcriptomic regulation^15^. Unlike CRISPRi—which uses DNA-targeting Cas9—the Cas13 is an RNA-targeting CRISPR-Cas effector capable of directly degrading RNA targets that are complementary to the spacer sequence of the gRNA^8,9,18^. Compared with RNAi, Cas13-based systems can achieve higher silencing efficiency, sequence specificity, and tolerance to RNA secondary structures^19–21^. Therefore, they have become a useful tool for elucidating the functional role of transcripts at the isoform level.

New developments in tools to modulate transcript expression have allowed for increased precision and potency in controlling the expression of the genes of interest. Individually, each tool offers a distinct trade-off between isoform resolution, potency, and potential off-target effects^13,19,20^. However, by selecting appropriate tools based on the desired goal of the research, they become powerful means of unambiguously determining the functionalities of transcripts stemming from the gene of interest. Overall, this report aims to present a framework for using CRISPRi and CRISPR-Cas13-mediated RNA degradation of the comprehensive discovery and elucidation of target transcript functional dynamics in cells. Furthermore, this study investigates the potential of the Cas13 system to target and silence specific transcript isoforms, including alternatively spliced mRNAs and circular RNAs (circRNAs). It also assesses the precision and efficacy of CRISPRi-mediated silencing and considers the limitations of using silencing outcomes to infer gene function.

## RESULTS

### Construction and validation of stable cell lines with inducible Cas expression

To ensure consistent expression of the desired Cas protein, a stable cell line was constructed by knock-in (KI) of the Cas protein cassette sequence into the T2 site of the *AAVS1* locus, according to previously reported methods. Subsequently, a cell line was established from single-cell colonies with the highest levels of reporter protein expression. The human PSC line WTB6 was used for the above experiments, whereas 1B4 (CRISPRi Gen 1 clone 4 derived from WTB6) was used as the reference cell line with a single KI at the *AAVS1* site (Fig. 1A). Briefly, the relevant sequence of the desired Cas protein and mCherry reporter was inserted into a vector containing the rtTA3G/TRE3G system for doxycycline (Dox) expression^17^. For the Cas proteins, HA-tagged ZIM3-dCas9 and RfxCas13d (CasRx) were used for CRISPRi and Cas13 silencing experiments, respectively, because of their reported advantages in potency and selectivity compared to other variants^15,19,22–24^.

**Figure 1:**
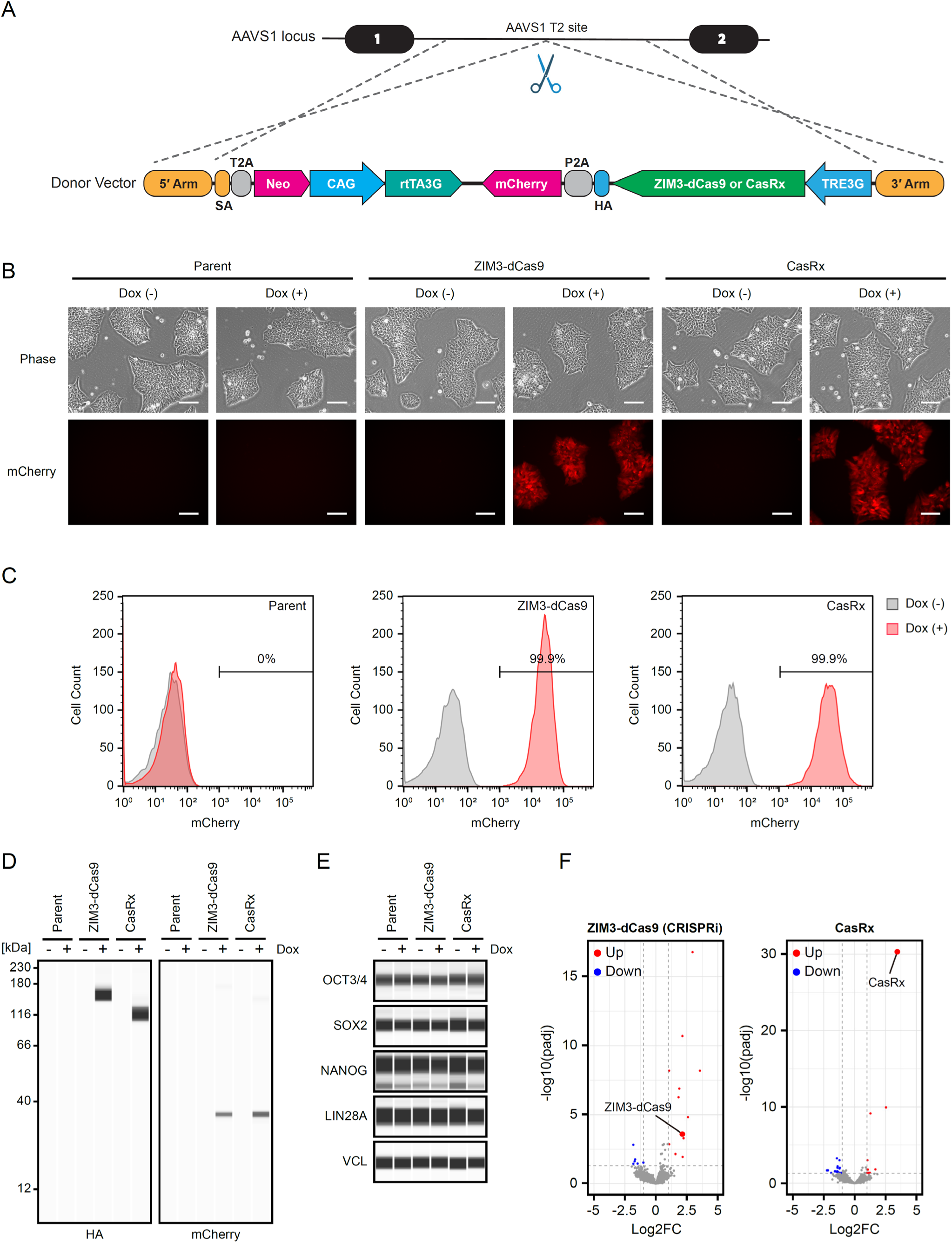
Generation of human PSC lines carrying inducible Cas. (A) Schematic of Dox-inducible Cas knock-in into the *AAVS1* allele. (B) Representative images of parental, ZIM3-dCas9, and CasRx cell lines in the presence or absence of Dox. Scale bars: 100 μm. (C) Representative flow cytometry results of mCherry expression in the cells depicted in (B). (D) Protein expression of HA-tagged Cas and mCherry in the cells shown in (B). (E) Protein expression of pluripotency markers in the cells depicted in (B). VINCULIN (VCL) was used as a loading control. (F) Volcano plot of RNA-seq analysis results following induction of the respective Cas proteins by Dox. Up-regulated and down-regulated genes are denoted in red and blue respectively. (|log_2_FC| > 1, adjusted *p* < 0.05).

Successful KI of the desired cassette was confirmed via genotyping PCR (Fig. S1A, B). For the KI allele-specific primer pairs, an amplicon of the expected length was observed in all KI cell lines, whereas no amplification was observed in the parental cell line. Conversely, amplification was observed only in the parental cell line and not in the KI cell line using wild-type (WT) allele-specific primers. The 1B4 line exhibited amplification with both the KI- and WT-specific primer pairs, and no amplification was observed in the absence of the template. Additionally, a copy number assay was performed using droplet digital PCR (ddPCR) and a probe targeting the neomycin resistance gene to assess the number of KI alleles in the genome. The 1B4 line was used as a reference with one copy of the KI allele per genome. The results indicated that the parental line had zero copies, whereas all KI lines had two copies of the KI alleles in their genomes (Fig. S1C). Finally, G-banding assessment of chromatin from the KI cell lines suggested that the KI of the cassette allele did not cause chromatin instability or rearrangement (Fig. S1D).

Thereafter, the efficiency of Dox-dependent induction of Cas expression by ZIM3-dCas9 and CasRx was evaluated. First, the expression of the mCherry reporter protein in the presence or absence of Dox was analyzed. In the presence of Dox, the KI cell line demonstrated a marked increase in mCherry fluorescence intensity compared with the parent cell line, which showed no fluorescence (Fig. 1B). This was further confirmed by flow cytometry, which showed that none of the cells in the parental cell line showed any change in fluorescence, regardless of the presence of Dox. Conversely, all KI cell lines exhibited increased fluorescence intensity following Dox treatment compared to the untreated cells (Fig. 1C).

Moreover, the expression of Cas and mCherry reporters was directly determined via western blotting using HA-tag and mCherry antibodies, respectively (Fig. 1D). The results revealed the induced expression of the respective Cas in KI cell lines, ZIM3-dCas9 (∼165 kDa) and CasRx (∼113 kDa) following Dox treatment. Conversely, no protein expression was observed in untreated cells. Additionally, the mCherry expression pattern was similar to that of the corresponding Cas protein. No protein was detected in the parental cell line, regardless of the presence of Dox.

Finally, this study assessed the global effects of KI and Cas expression on cellular identity and transcriptomic changes. To this end, the expression of the protein markers associated with pluripotency, OCT3/4, SOX2, NANOG, and LIN28A, was analyzed (Fig. 1E)^25^. The results of western blotting indicated consistent expression of the selected markers across all tested cell lines, regardless of the cassette sequence KI or expression of Cas resulting from Dox treatment. Additional changes in the transcriptomic environment were assessed using RNA-seq (Fig. 1F). Differential analysis of gene expression before and after the induction of Cas expression revealed a significant upregulation of transcripts from the inserted vector (ZIM3-dCas9-P2A-mCherry and CasRx-P2A-mCherry, respectively), with only minimal perturbation of the expression of endogenous transcripts.

### CRISPRi and CasRx achieve comparably high KD efficiency

To evaluate the efficacy of the prepared cells for KD experiments, the relevant sgRNA was designed using available bioinformatic algorithms^26–28^. Its stable expression was achieved by insertion via the piggyBac transposon vector system, as reported previously^29^. Following the introduction of Dox into the culture media, the expression of Cas protein was induced, and the cells were subsequently analyzed after 5 d of Dox treatment. Here, a typical marker expressed in undifferentiated PSCs—*POU5F1* encoding transcription factor OCT4—was targeted for KD. Thereafter, changes in the gene and protein expression were analyzed via RT-qPCR and immunocytochemistry (ICC). Following Dox treatment, CRISPRi and CasRx achieved a 99.4% (p < 0.0001) and 95.5% (p < 0.0001) reduction in *POU5F1* expression, respectively, compared to the cells expressing non-targeted (NT) sgRNA (Fig. 2A). This was further reflected by ICC analysis of OCT4 protein expression in KD cells, which demonstrated the abolishment of OCT4 protein signals upon Dox treatment. In contrast, cells expressing control non-targeting sgRNA (sgNT) did not exhibit any changes in the expression of their respective genes (Fig. S2A, B). Moreover, to confirm the functional loss of OCT4, the expression levels of the downstream genes regulated by OCT4 —*NANOG*— was determined. In both cases, an 88.9% (p < 0.0001) and 82.9% (p < 0.0001) reduction in the expression of *NANOG* was observed, respectively for CRISPRi and CasRx, compared to the NT cells (Fig. 2B).

**Figure 2:**
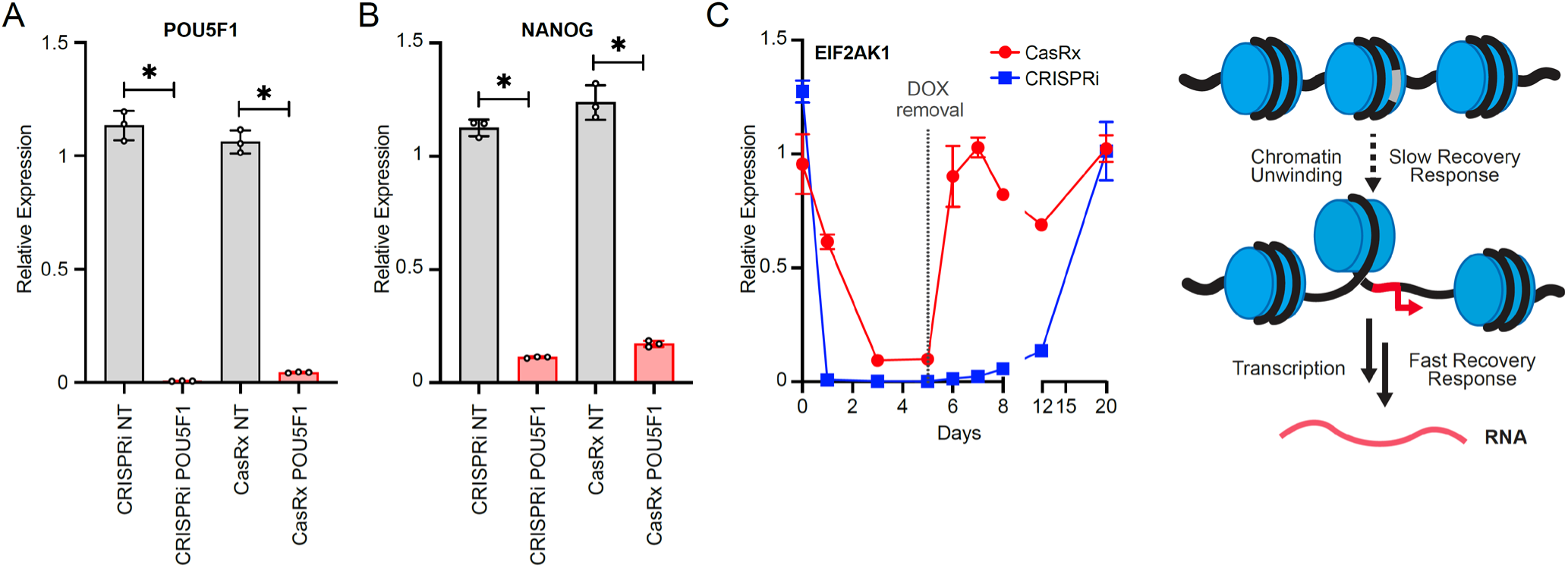
Comparison of knockdown behavior between CRISPRi and CasRx. (A) Relative expression of PSC marker *POU5F1* following KD using CRISPRi and CasRx (mean ± SD, n = 3, *: *p*_CRISPRi_ = 0.0011, *p*_CasRx_ = 0.00081, Student’s t-test). (B) Relative expression of *NANOG* in (A) (mean ± SD, n = 3, *: *p*_CRISPRi_ = 0.00039, *p*_CasRx_ = 0.0013, Student’s t-test). (C) Time-dependent analysis of *EIF2AK1* KD and its recovery (left), along with a schematic representation of the underlying mechanisms resulting in the difference in gene recovery speed (right).

### CRISPRi and CasRx differ in silencing and reversal behavior

The primary benefit of using non-destructive CRISPRi and CasRx lies in its ability to reverse gene silencing, wherein withdrawal of Dox abrogates the Cas effector protein and permits the restoration of target gene expression^17^. To evaluate the gene silencing characteristics associated with CRISPRi and CasRx, a time-course analysis of gene silencing was performed. Silencing of the non-essential gene, *EIF2AK1*,^26^ by CRISPRi and CasRx was induced for 5 d before the withdrawal of Dox, and the recovery of the target gene was tracked for up to 20 d (Fig. 2C, left). CRISPRi achieved >99.9% repression of the target gene on day 1 following Dox treatment and plateaued until Dox removal on day 5. Following the removal of Dox and the abolishment of the expression of the ZIM3-dCas9 effector protein, complete recovery of silenced gene expression was achieved after day 20 (Fig. 2C, right). Conversely, the KD of CasRx achieved a lower maximal (>90%) repression of the target gene by day 3, with a complete recovery of gene expression 1 d after withdrawal of Dox on day 6.

### KD characteristics of CRISPRi and CasRx

#### Limitation of CRISPRi

Despite the potent KD demonstrated by CRISPRi, the mechanism underlying transcriptomic repression introduces potential limitations to the range of targetable genes. Genes with a transcription start site (TSS) in proximity to or overlapping each other could potentially be knocked down in tandem, resulting in unwanted off-target effects and misinterpretation of the KD gene phenotype (Fig. 3A).

**Figure 3:**
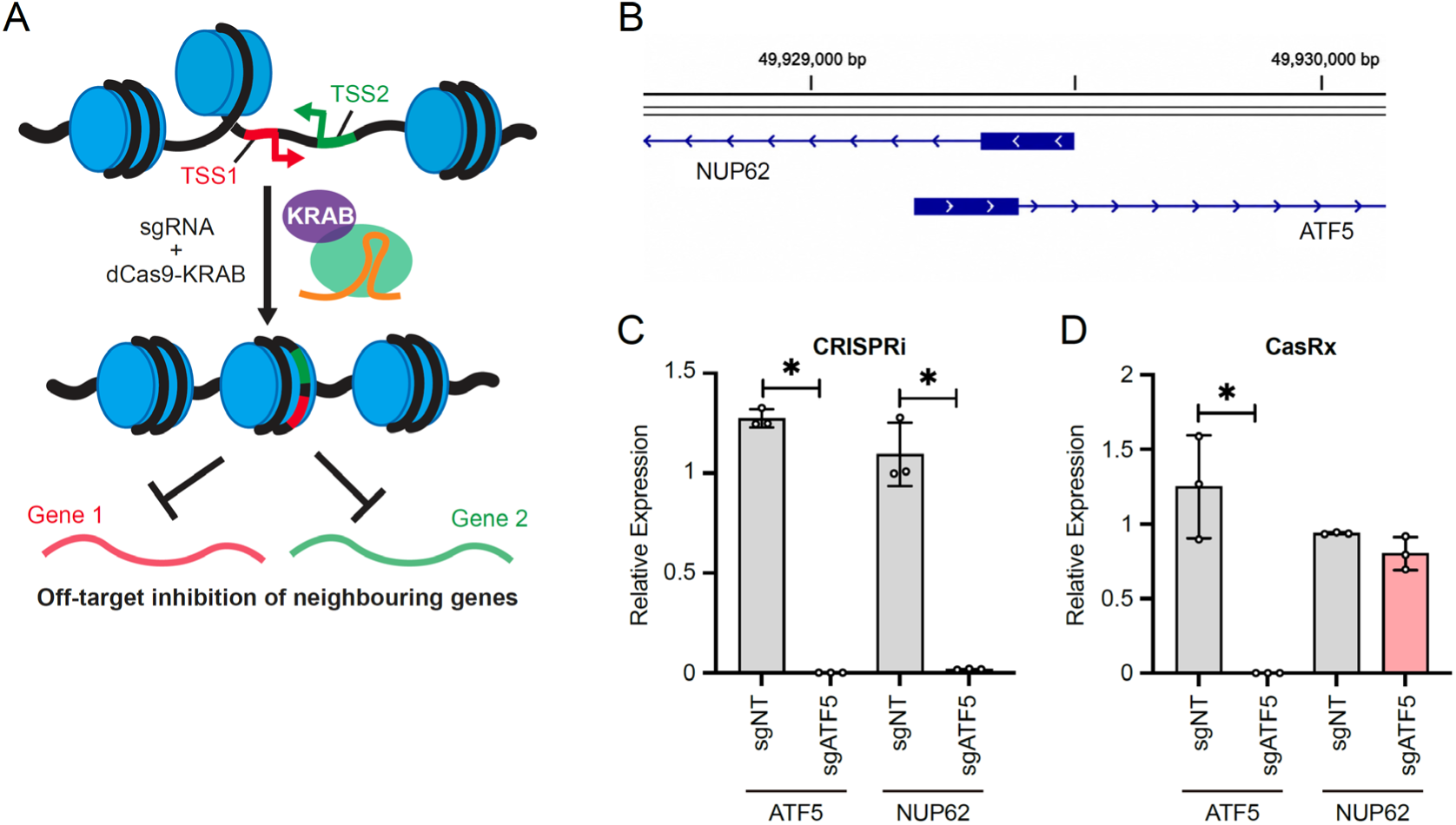
A model for overcoming the limitations of CRISPRi using CasRx. (A) Schematic representation of the off-target mechanism of CRISPRi. (B) Genomic location and overlap between *ATF5* and *NUP62*. Relative expression of *ATF5* and *NUP62* following sgATF5 dependent KD using (C) CRISPRi or (D) CasRx (mean ± SD, n = 3, CRISPRi *: *p*_ATF5_ = 0.00043, *p*_NUP62_= 0.0072; CasRx *: *p*_ATF5_ = 0.024, Student’s t-test).

We previously conducted a whole-transcriptome CRISPRi KD screening assay to elucidate the essential genes for PSC survival^30^. Analysis of the screening results indicated that a transcription factor, ATF5, was one of the top essential genes, with a potent phenotype upon KD. This was confirmed by a follow-up gene-specific CRISPRi experiment, which resulted in significant cell death upon *ATF5* KD, as visualized by 7-Amino-Actinomycin D (7-AAD) staining (Fig. S3A, top). However, further analysis of the gene locus revealed that *ATF5* shares an overlapping TSS with *NUP62*, indicating a potential off-target effect of CRISPRi on the latter (Fig. 3B)^31^. A follow-up gene expression analysis of CRISPRi *ATF5* KD cells was conducted to assess the expression of *NUP62*. The results showed *NUP62* expression KD by 99.3% (p < 0.0001), in addition to that of the intended target *ATF5*, which showed KD of >99.9% (p < 0.0001) (Fig. 3C).

To clarify the origin of the potent KD phenotype observed with CRISPRi, further investigations were performed using orthogonal siRNA-mediated KD of *ATF5* and *NUP62*. siRNA KD of *ATF5* did not result in any observable change in cell death compared to the control (Fig. S3B, siNC vs siATF5). In contrast, siRNA KD of *NUP62* resulted in a lethal cell phenotype (Fig. S3B, siNUP62). Illustrating the presence of the *ATF5*-*NUP62* off-target gene pair in CRISPRi KD experiments. Considering these results, other potential off-target gene pairs were investigated and retrieved from publicly available gene annotations. The analysis is provided in Table S1.

#### CasRx allows selective KD in transcript- and isoform-specific manner

As the target molecule of CasRx is RNA, the entire transcriptomic sequence could theoretically be exploited to achieve KD. This provides a larger pool of potential sgRNA designs and allows for superior target-specificity.

This was demonstrated by performing KD of *ATF5* using CasRx and assessing the off-target effects observed in the CRISPRi experiment. The results revealed a KD efficiency of the *ATF5* gene to that of CRISPRi, with >99.9% (p < 0.0001) reduction in expression without the previously observed off-target effect on *NUP62* (Fig. 3D, Fig. S3A, bottom).

Similarly, the selective KD of genes are traditionally non-targetable by CRISPRi, owing to their sequence homogeneity in the TSS, was demonstrated. For instance, *ESRG,* which uses a human endogenous retrovirus-H (HERV-H) LTR7 for its TSS^32^, was targeted for KD using an sgRNA-targeting an internal exon (Fig. 4B). The results showed an 88.7% KD (p < 0.005) of *ESRG* without affecting the expression of the other genes—including *RPL38L*, *IL34*, and *KLKB1*—that share the same TSS sequence.

**Figure 4:**
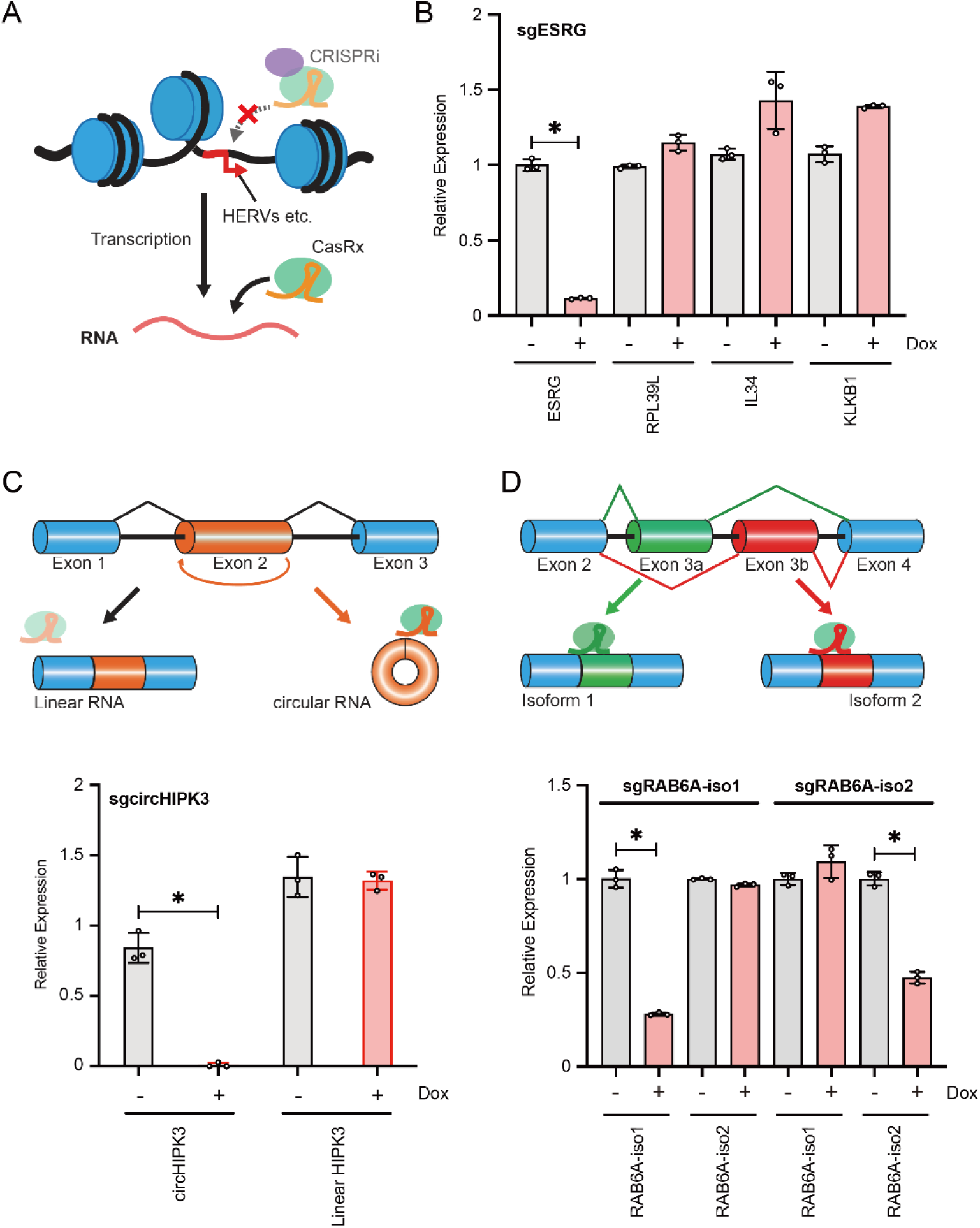
Knockdown selectivity and sgRNA design for CasRx. (A) Schematic representation of CasRx-targeting genes that are untargetable using CRISPRi. (B) Relative expression of ESRG and other genes sharing the same HERV-H LTR7 TSS following CasRx KD (mean ± SD, n = 3, *: *p*_ESRG_ = 0.00047, Student’s t-test). (C) Schematic representation and selective KD of circHIPK3 vs linear HIPK3 mRNA (structural isoform) (mean ± SD, n = 3, *: *p*_circHIPK3_ = 0.0046, Student’s t-test). (D) Schematic representation and selective KD of RAB6A-iso1 and RAB6A-iso2 (splice isoform) (mean ± SD, n = 3, *: *: *p*_RAB6A-iso1/sgRAB6A-iso1_ = 0.0011, *p*_RAB6A-iso2/sgRAB6A-iso2_ = 0.000051, Student’s t-test).

Moreover, the target specificity of CasRx can be extended to differentiate between targets beyond the gene and transcript levels. Alternatively spliced transcript isoforms can be specifically targeted for KD through the design of sgRNAs spanning splice junctions uniquely associated with the spliced isoform in question, without affecting the expression of other variants.

Exon 2 of the *HIPK3* gene reportedly undergoes a special form of alternative splicing—known as back-splicing—to produce a distinct circular RNA isoform from the second exon of the *HIPK3* gene, which is identified as circHIPK3 (hsa_circ_0000284)^33^. By designing sgRNA to specifically overlap across the back-spliced junction sequence^34,35^, CasRx was able to achieve >99.9% KD (p < 0.0001) of the circHIPK3 isoform without affecting the expression of its linear counterpart (Fig. 4C). Similarly, isoforms derived from genes with mutually exclusive exons can be identified and targeted. The *RAB6A* gene contains exons 3a and 3b, which are mutually exclusive and give rise to two distinct splicing isoforms, RAB6A-iso1 and RAB6A-iso2, respectively (Fig. 4D)^36^. SgRNA designed to overlap the specific splice junctions resulting from mutually exclusive splicing were able to achieve specific KD in the expression of only the targeted isoform without affecting the expression of its counterpart, albeit with varying efficiencies (76% vs. 53% KD for RAB6A-iso1 and RAB6A-iso2, respectively). However, it must be noted that the limited design window for sgRNAs and potential competition from pre-attached splicing complexes may cause failure in the KD of the desired splice isoform by CasRx. This was especially evident for circRNA targets, in which the major circRNA isoforms stemming from the genes *FAT1* (circFAT1, hsa_circ_0001461), *AMBRA1* (circAMBRA1, hsa_circ_0007001), and *CCDC134* (circCCDC134, hsa_circ_0001238) were resistant to CasRx-mediated degradation (Supporting Fig. S4).^35^

## DISCUSSION

Achieving robust and interpretable loss-of-function phenotypes requires gene knockdown experiments under standardized conditions. Therefore, choosing an appropriate knockdown methodology and streamlining the preparation of KD-competent cell lines are essential for reducing technical noise and increasing reproducibility across experiments and laboratories. This study presents a systematic method that allows for the consistent preparation of cell lines capable of inducing the expression of the chosen Cas effector protein used for the two most prevalent Cas-dependent knockdown methods, CRISPRi and CasRx, without disrupting genome stability or the expression of the host transcriptomic environment. Both methods have similarly robust KD efficiencies, resulting in the loss of the target protein and changes in downstream gene expression, supporting their application in targeted loss-of-function analyses. Finally, a thorough time-course analysis and comparative studies were conducted to elucidate the KD behavior of the two methods.

CRISPRi had a measurable advantage over CasRx in terms of KD potency across all genes tested, achieving KO-level downregulation. Furthermore, CRISPRi enables a more rapid KD of the target gene but requires an extended time for recovery owing to slow gene reactivation through chromatin unwinding. However, despite the potency of this methodology, targeting the TSS has inherent limitations resulting from the narrow window of targetable sequences and the long-range effects of chromatin remodeling. To avoid potential off-target effects, such as those observed for the *ATF5*-*NUP62* gene pair, the gene of interest should be vetted against the gene pair list and known retrotransposon TSSs before performing KD experiments. Additionally, the expression of neighboring genes should be rigorously tested after KD is achieved.

Conversely, CasRx has distinct advantages in terms of specificity owing to the large sequence window available for the design of sgRNAs, providing better precision for KD and allowing specific splice isoform targeting. This is advantageous when attempting to target families of genes that use TSS stemming from retrotransposable elements (Fig. 4A), such as the HERV family and mammalian apparent LTRs (MaLRs)^37^. Compared with CRISPRi, CasRx demonstrated slower kinetics in achieving maximal KD efficiency, as it competes with the speed of the transcription machinery while allowing for faster recovery when the nuclease is removed. However, despite its versatility and the availability of state-of-the-art prediction algorithms for sgRNA design, KD efficiency deviates across different RNA targets and isoforms, with some being completely resistant to degradation. This is especially problematic in cases where the window of targetable sequences is restricted to discriminate between splice isoforms, which limits the diversity of designable sgRNAs and the likelihood of achieving a satisfactory KD.

Overall, CRISPRi and CasRx are robust methods for achieving KD of target genes. Nonetheless, this study highlighted the distinct advantages and limitations that should be considered when selecting a KD strategy. This study provides insights into the unreported pitfalls and strategies to avoid them, thus providing a valuable resource for future experimental designs. Furthermore, despite their apparent competing utilitarianism, we believe that the two methodologies should be used in a complementary manner. The rapid and leakless KD of CRISPRi was used for the initial analysis of gene functionality, followed by the use of CasRx as an orthogonal method to clarify the presence of off-target effects and determine the specific isoform associated with the observed gene function.

### Limitations of the study

This study established a standardized, inducible human PSC platform for high-efficiency knockdown using CRISPRi (ZIM3-dCas9) and CasRx (RfxCas13d). Direct comparisons of modality-specific behaviors (e.g., potency, reversibility, and isoform-level targeting) were conducted to provide general guidance for tool selection. The primary limitation of CRISPRi is the interpretability of the KD phenotype, owing to the complex promoter architecture of the genome. This is illustrated by *ATF5*-*NUP62* co-suppression due to their overlapping TSS, resulting in confounded phenotypes. Therefore, additional care must be taken to validate the TSS positions of neighboring genes to avoid misattributions. CasRx offers orthogonal RNA-level targeting, which addresses the limitations of TSS targeting and allows isoform-specific targeting. However, knockdown generally proceeds more slowly and often reaches lower maximal repression than CRISPRi. Additionally, KD efficacy can vary substantially depending on the target, with some RNAs exhibiting varying degrees of resistance to CasRx degradation. As such, isoform-specific applications are not uniformly reliable in practice because junction targeting can be constrained by narrow design windows and interference from RNA-processing complexes. Targeting circRNAs is particularly challenging because several circRNAs are seemingly immune to CasRx degradation.

## ACKNOWLEDGMENTS

We thank B. Conklin, Y. Sato, M. Taipale, K. Woltjen, S. Yamanaka, F. Zhang, and CiRA Common Equipment Management Office for sharing materials and equipment, A. Ogawa, for technical assistance, Y. Ishida and S. Takeshima for administrative support, and Editage for English language editing. The schematic figures were created in BioRender.com.

This work was supported by Grants-in-Aid for Scientific Research from the Japanese Society for the Promotion of Science (JSPS) 20K20585 (KT), Grants-in-Aid for Scientific Research from JSPS 24H00568 (KT), Grants-in-Aid for Scientific Research from JSPS 25K18461 (CO), Grants-in-Aid for Scientific Research from JSPS 25K18211 (LN), Core Center for Regenerative Medicine and Cell and Gene Therapy from Japan Agency for Medical Research and Development (AMED) JP23bm1323001 (KT), Core Center for iPS Cell Research from AMED JP21bm0104001 (KT), a grant from the Takeda Science Foundation (KT), a grant from the Uehara Memorial Foundation (KT), a grant from the Mochida Memorial Foundation (KT), a grant from the Stellar Science Foundation (LN), Support for Pioneering Research Initiated by the Next Generation from Japan Science and Technology Agency (JST) (TM, SS), and The iPS Cell Research Fund from the Center for iPS Cell Research and Application, Kyoto University (LN, CO, KT).

## AUTHOR CONTRIBUTIONS

Conceptualization, L.N. and K.T.; methodology, L.N., C.O., and K.T.; Investigation, L.N., T.M., S.S.,

H.M., M.N.,C.O., and K.T.; writing—original draft, L.N., T.M., and K.T.; writing—review & editing, L.N.,

T.M., C.O., and K.T.; funding acquisition, L.N., T.M., C.O., and K.T.; resources, L.N., T.M., and K.T.; supervision, L.N., and K.T.

---

**Figure S1:**
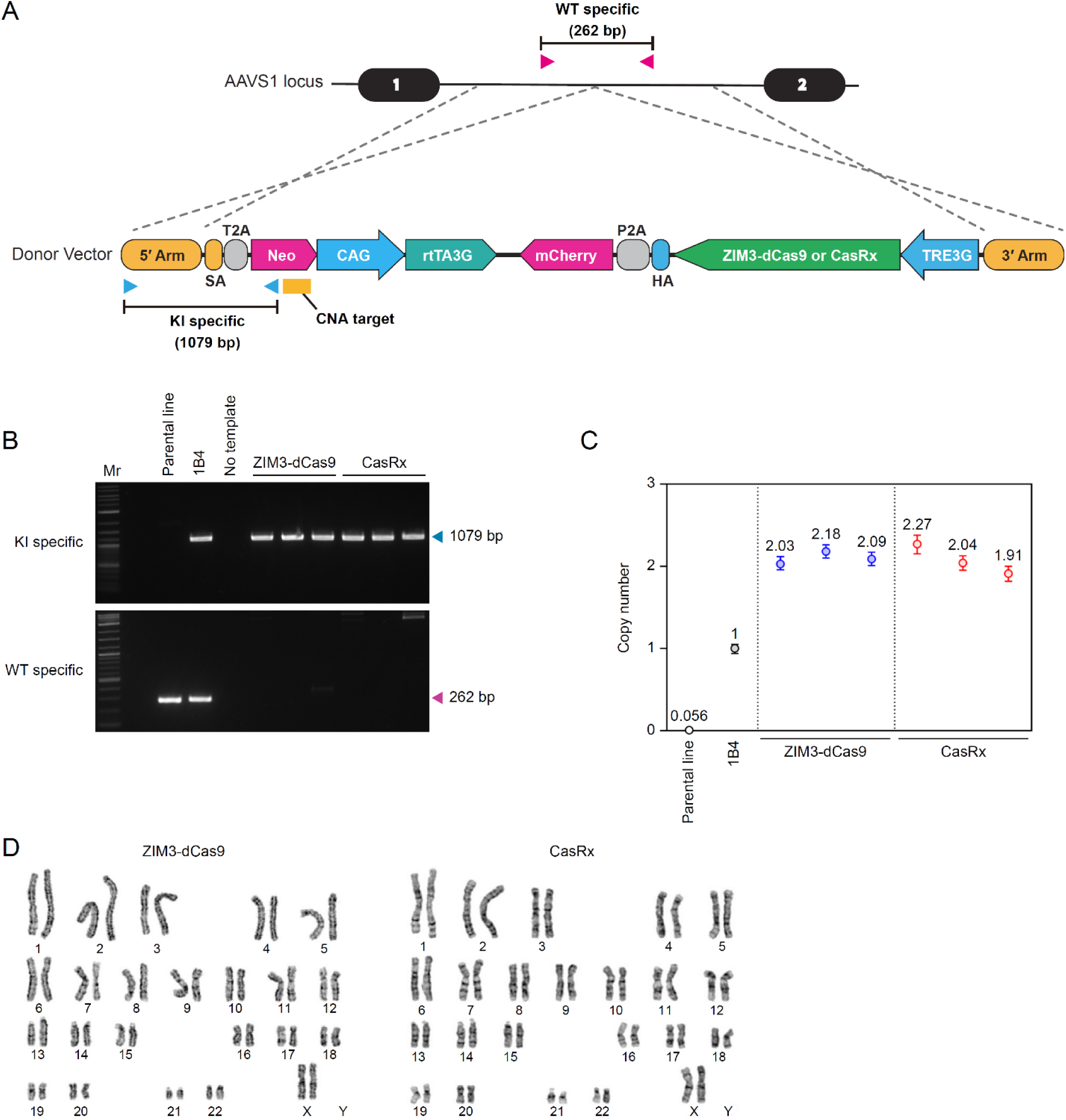
Genotyping of KI cell lines. (A) The panel showing the primer sets detecting Wild-type (WT, magenta) and Knock-in (KI, blue) allele, and the probe for copy number assay (CNA, yellow). (B) The genotyping PCR for KI (top) and WT (bottom) allele. (C) The copy number of KI alleles detected by ddPCR. (D) The G-banding shows the clones have no apparent karyotype abnormalities.

**Figure S2:**
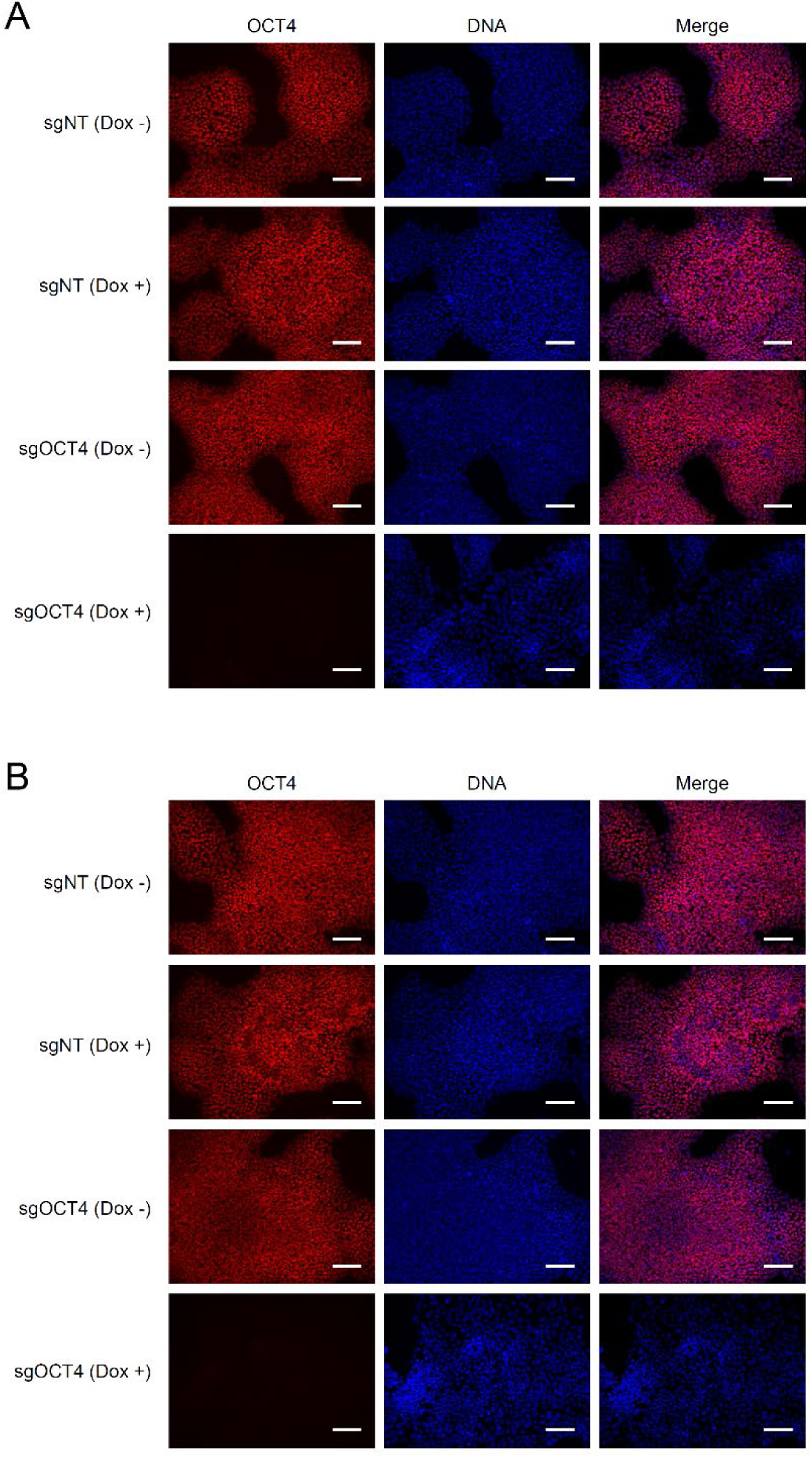
Protein expression of OCT4. ICC image comparing the expression of OCT4 protein following sgNT or sgOCT4 with (Dox +) or without (Dox -) Dox treatment for (A) CRISPRi and (B) CasRx dependent KD (right: OCT4, middle: DNA, left: merged). Scale bars: 100 μm.

**Figure S3:**
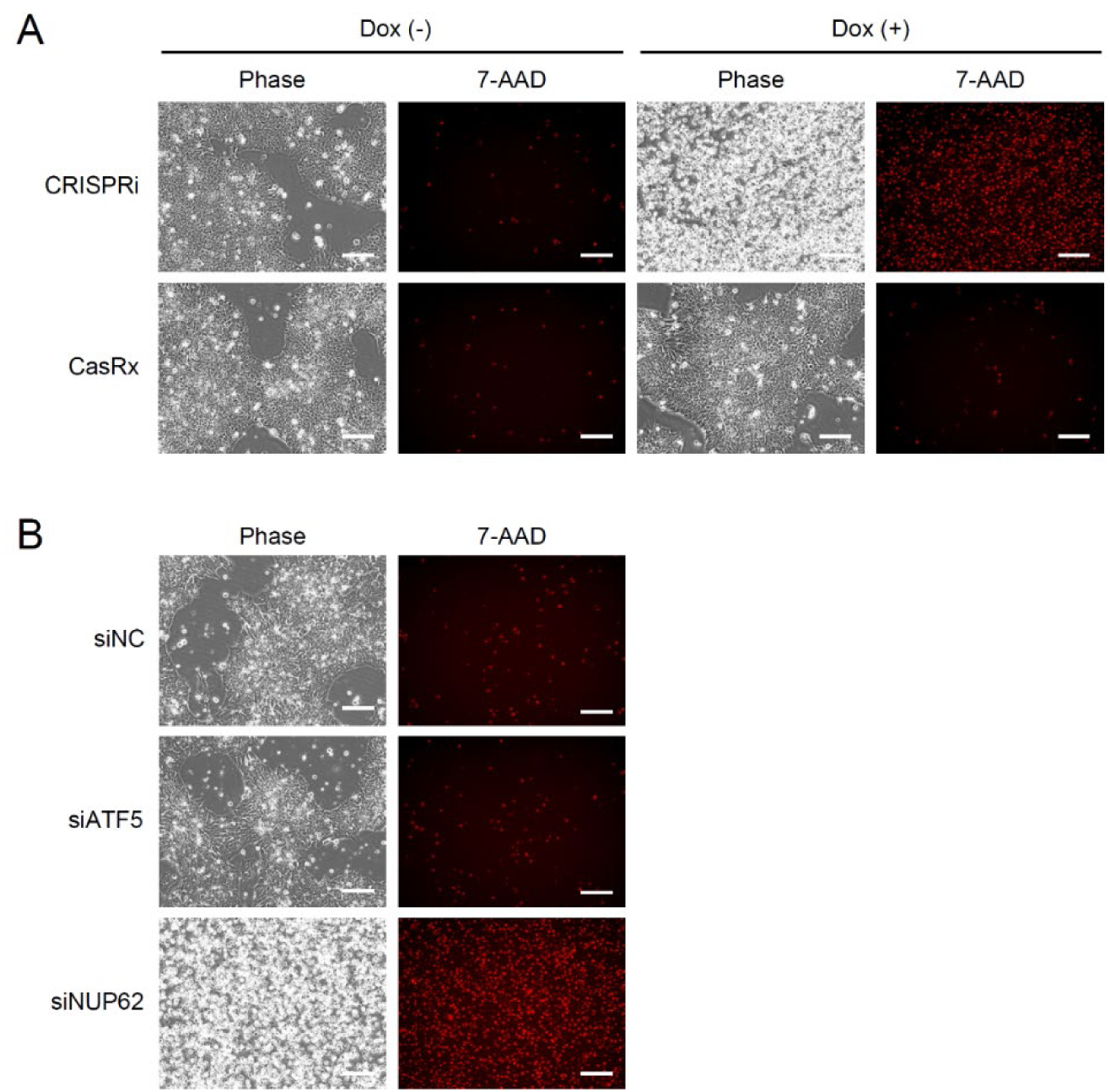
Off-target effect of ATF5 KD. Cell death upon (A) CRISPRi (top) or CasRx (bottom) KD of the ATF5 gene, and (B) siRNA dependent KD of ATF5 and NUP62. The cellular death was visualized by 7-AAD staining. Scale bars: 100 μm.

**Figure S4:**
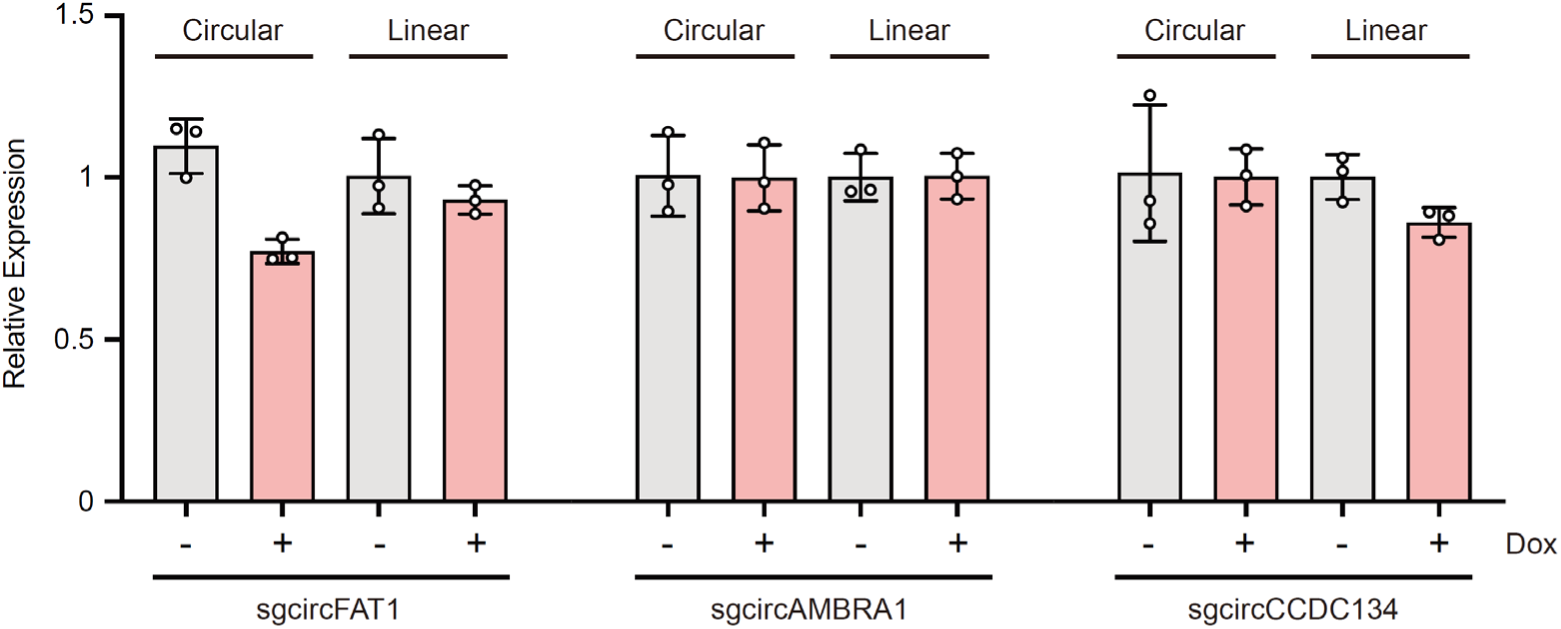
Limitation of circRNA KD. The graph shows examples of circRNA isoforms that resist degradation by CasRx (circFAT1, circAMBRA1, and circCCDC134, from left to right).

## KEY RESOURCES TABLE

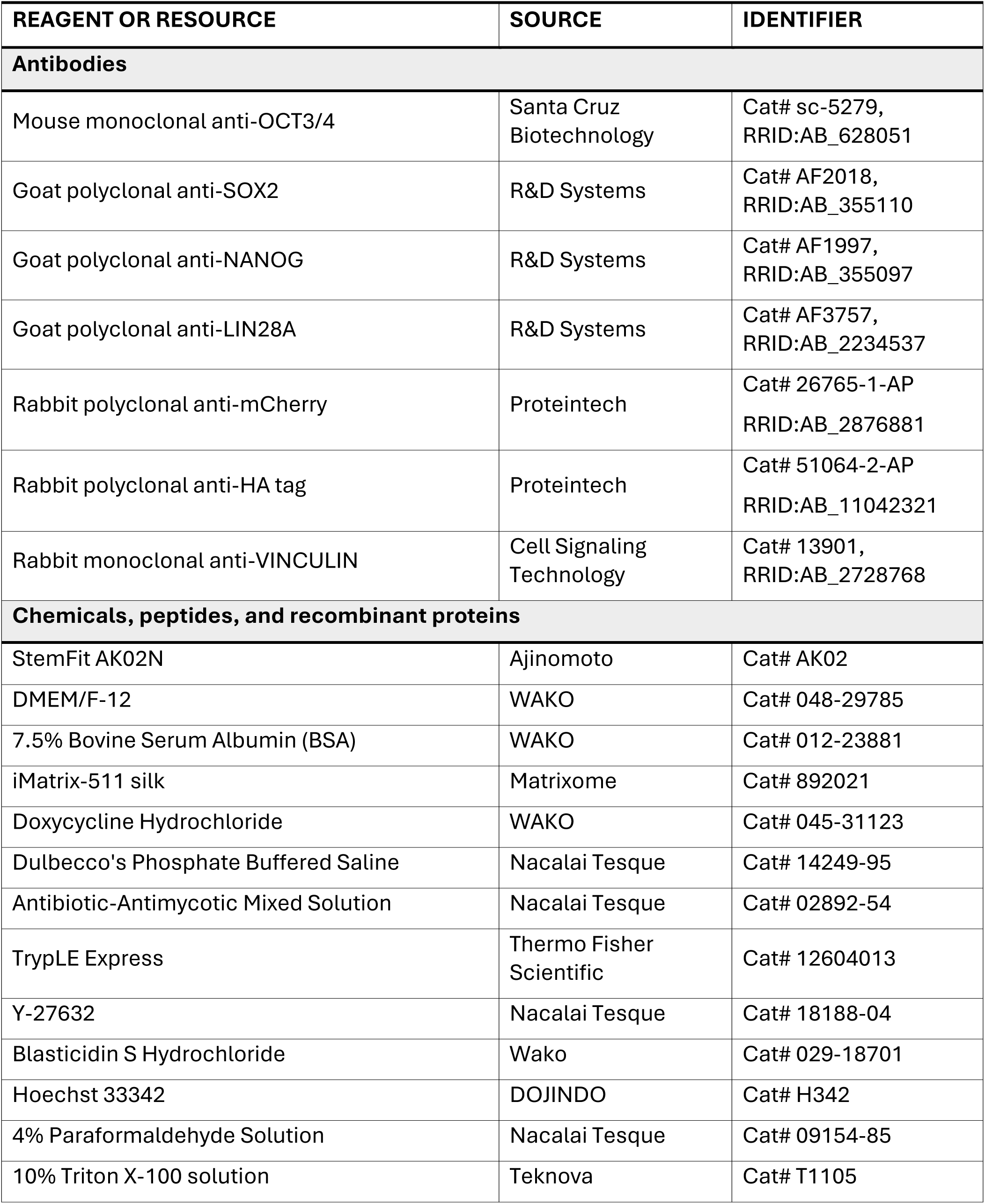
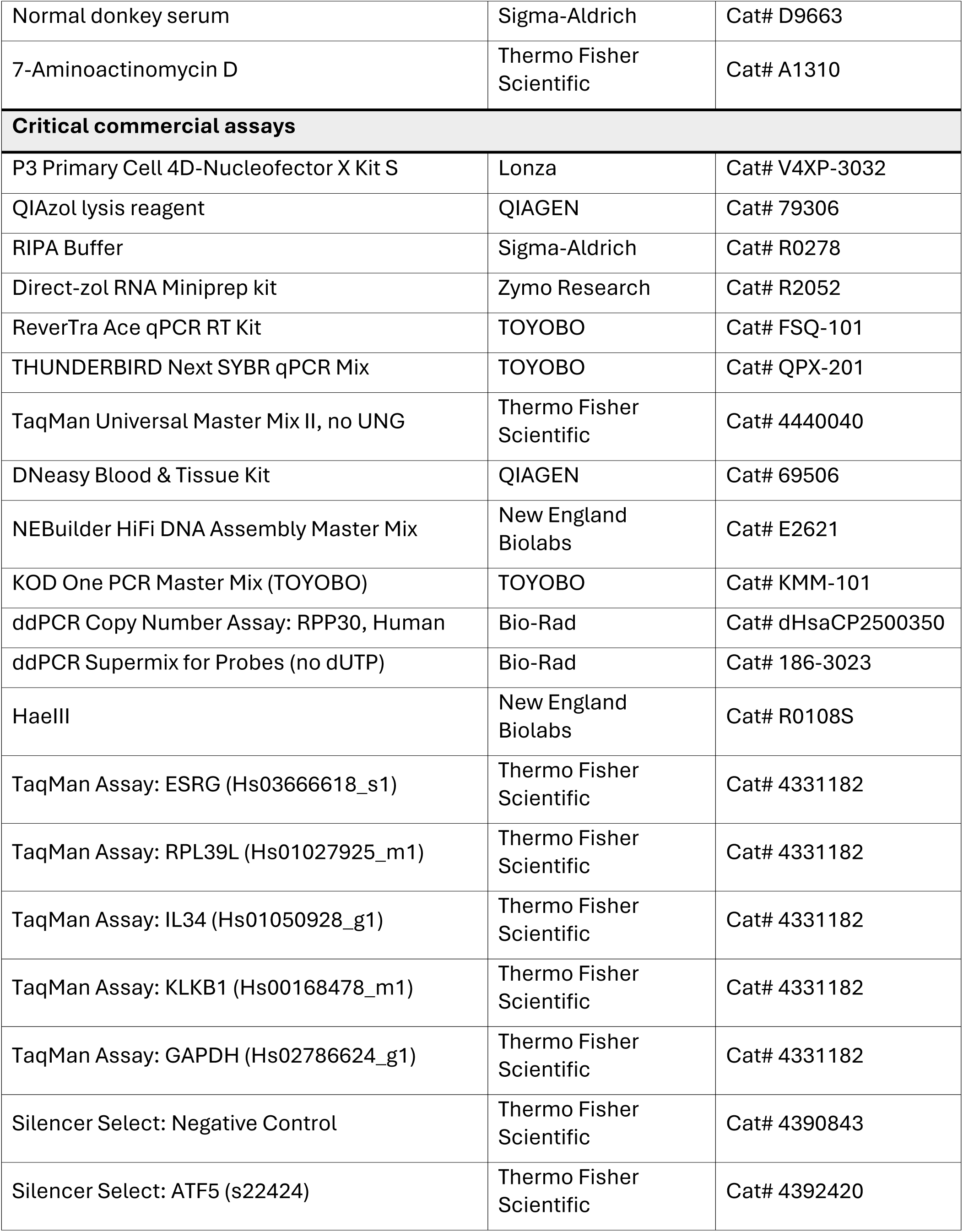
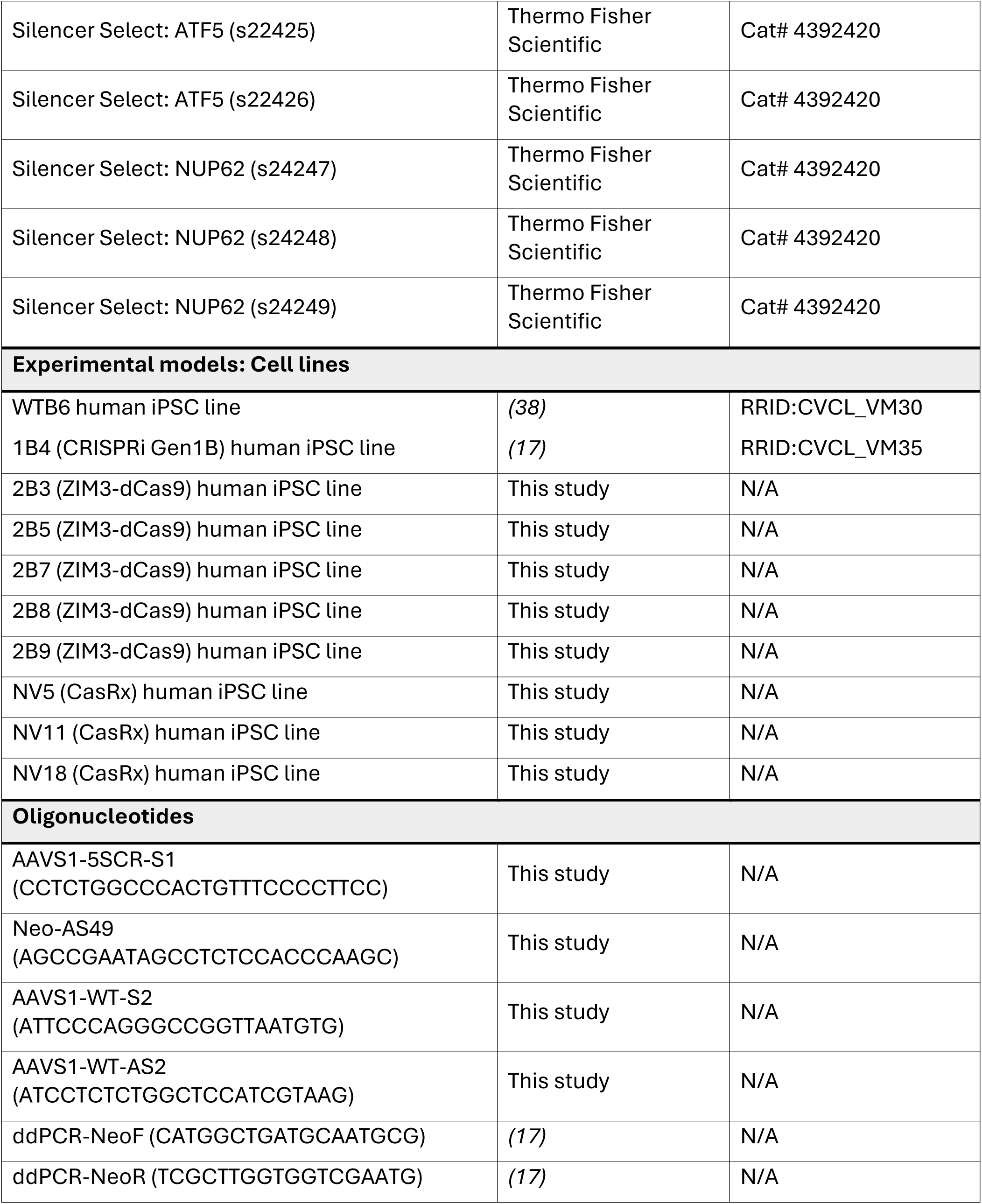
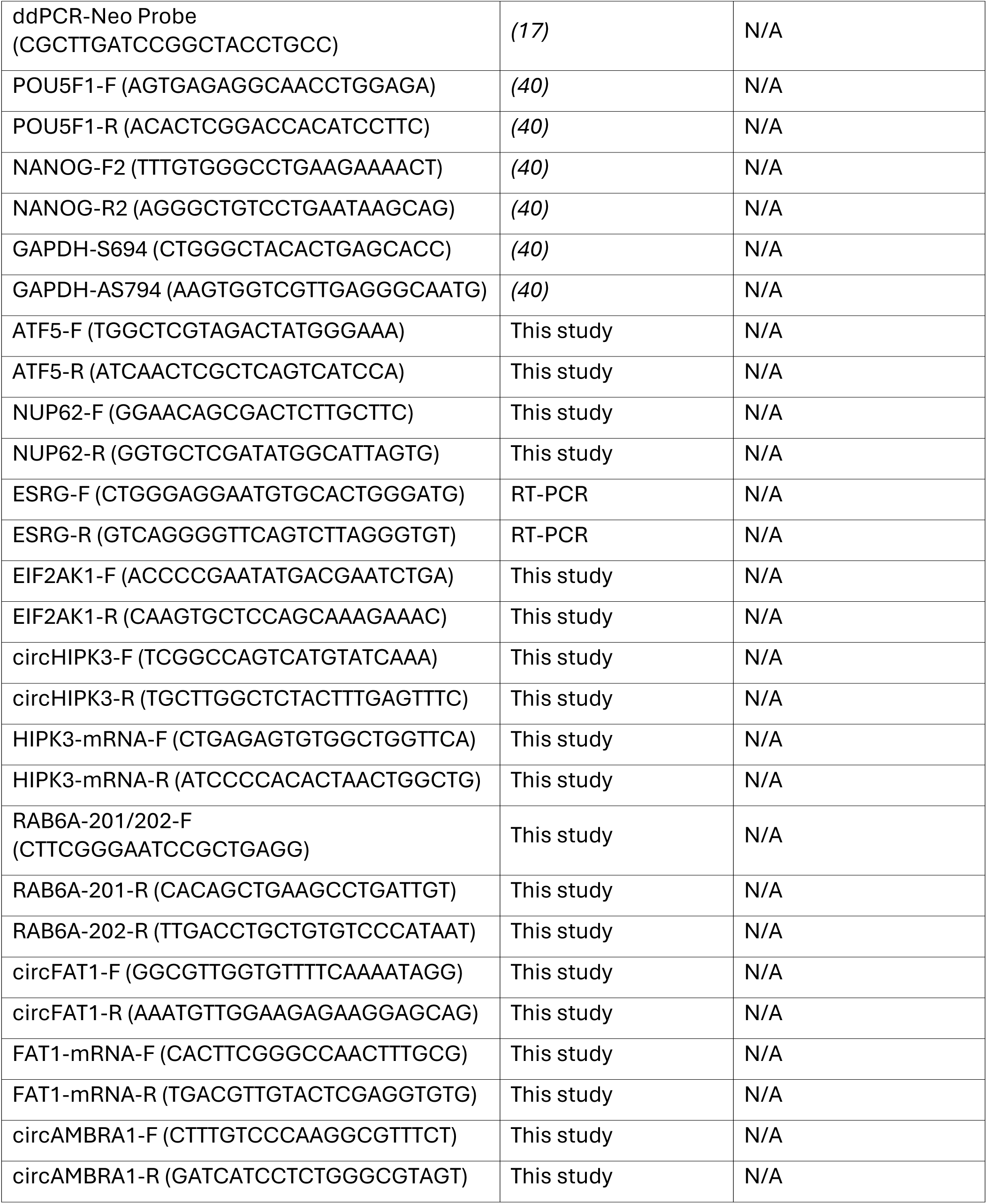
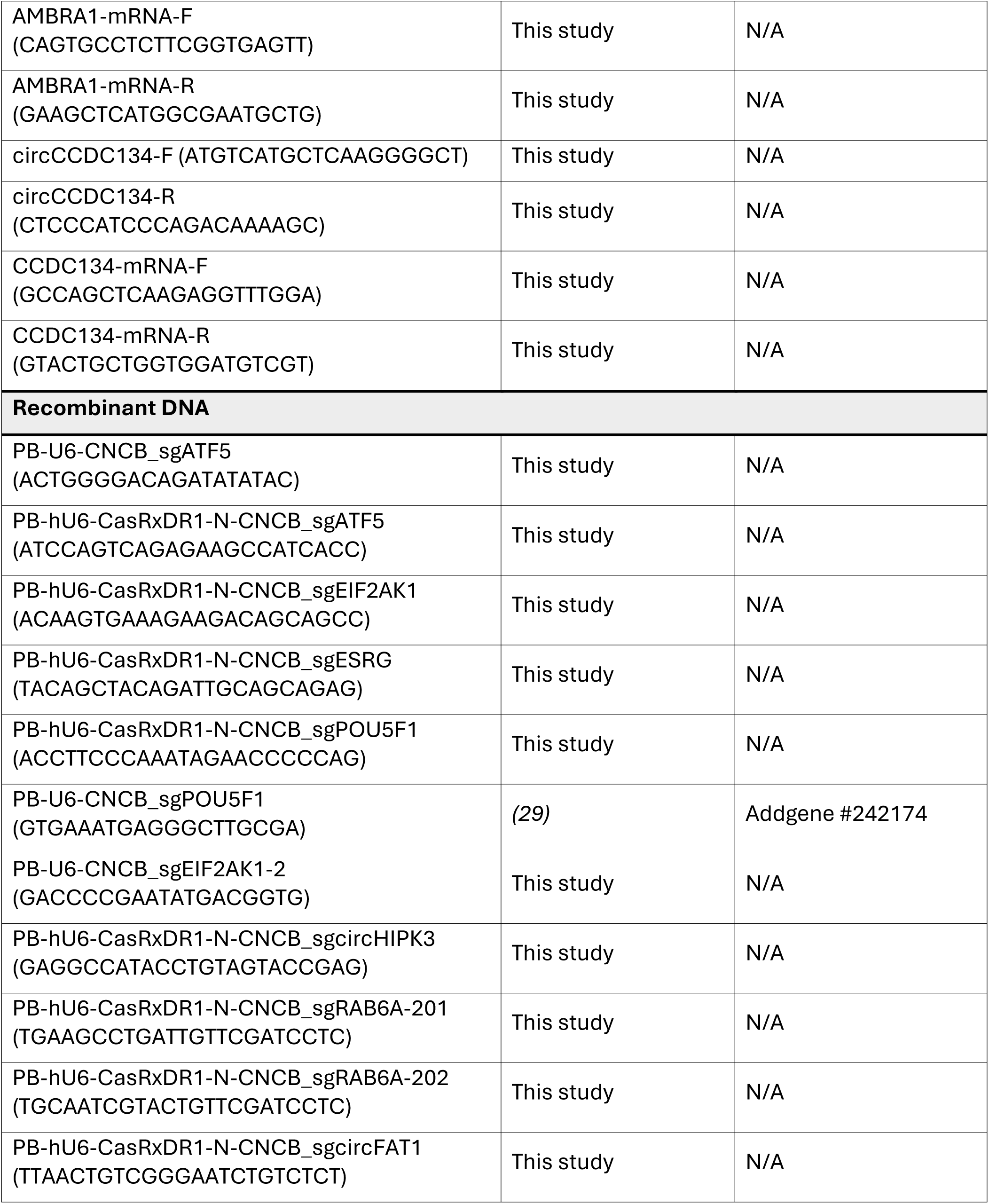
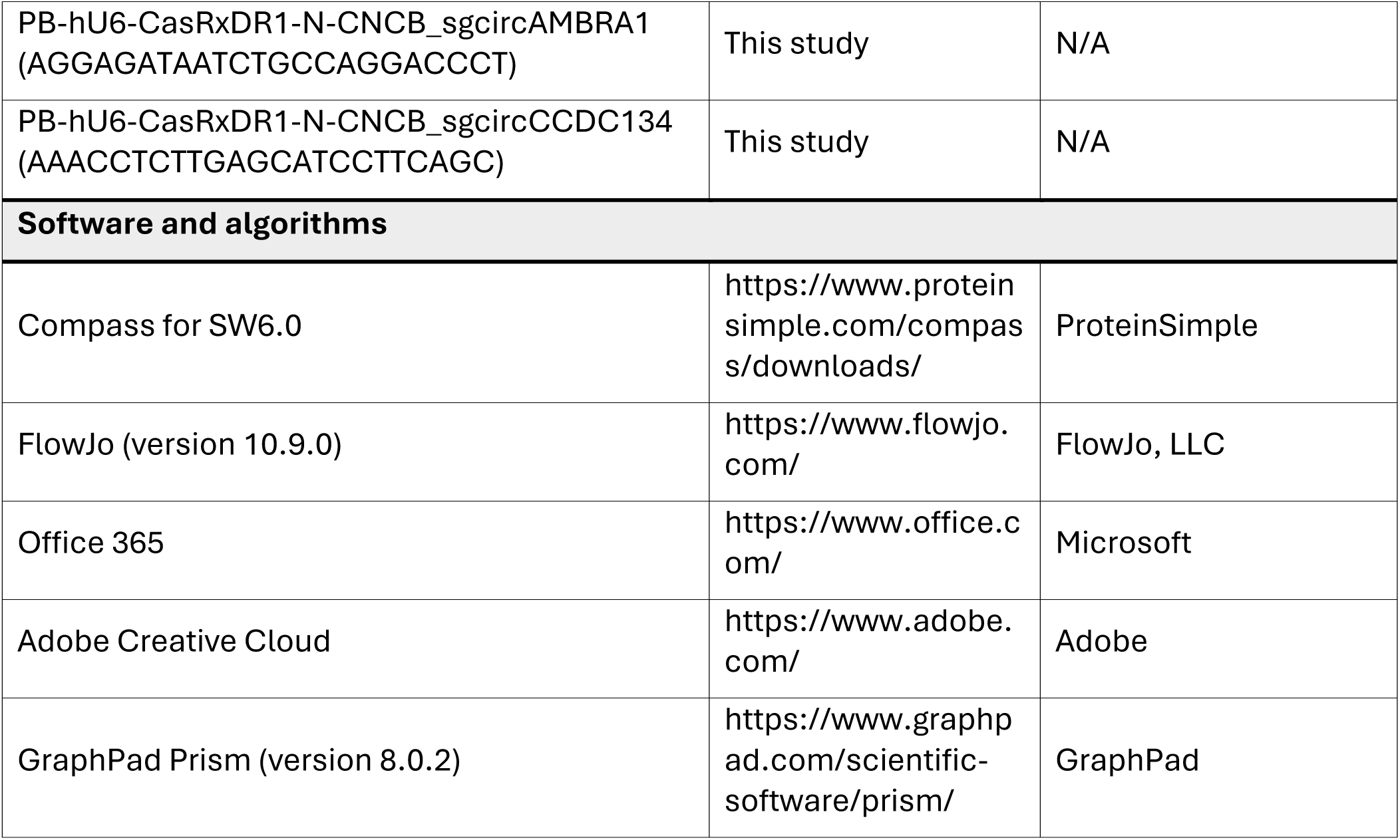

## EXPERIMENTAL MODEL AND STUDY PARTICIPANT DETAILS

Human induced pluripotent stem cell (iPSC) lines WTB6 (RRID:CVCL_VM30) and 1B4 (CRISPRi Gen 1 clone 4 derived from WTB6) (RRID:CVCL_VM35) were provided by Dr. Bruce R. Conklin (Gladstone Institutes and University of California, San Francisco, CA, USA)^17,38^. The cells were cultured in humidified incubators at 37°C in 5% CO_2_ and 20% O_2_. All reagents were warmed in a water bath set at 23°C before use, unless otherwise noted. Periodic tests were performed to confirm that all cell lines used in this study were negative for mycoplasma contamination. Karyotype analysis was performed by Nihon Gene Research Laboratories Inc.

## METHOD DETAILS

### Cell lines and culture conditions

Human PSCs were cultured in tissue culture plates coated with iMatrix-511 silk (Matrixome) and maintained in StemFit AK02N medium (Ajinomoto), as previously reported^40^. For routine passaging, the cells were rinsed once with Dulbecco’s phosphate-buffered saline (D-PBS; Nacalai Tesque) and treated with TrypLE Express (Thermo Fisher Scientific) for 10 min at 37°C. The cells were dissociated into single-cell suspensions and washed with Dulbecco’s Modified Eagle Medium/Ham’s F-12 (DMEM/F-12; Wako Chemicals) supplemented with 0.1% bovine serum albumin (BSA; Wako Chemicals). Following cell counting and centrifugation, the cells were resuspended in StemFit AK02N medium containing 1.67 μg/mL iMatrix-511 silk, and 10 μM Y-27632 (Nacalai Tesque)^41^. The karyotype integrity of all PSC lines used in this study was confirmed via G-banding analysis performed by Nihon Gene Laboratories, which revealed no chromosomal abnormalities.

### Plasmid construction

The ZIM3 KRAB used in this study was generated via DNA synthesis based on the sequence of pLX303-ZIM3-KRAB-dCas9 (Addgene Plasmid #154472: a gift from Mikko Taipale)^22^. Similarly, the CasRx sequence was synthesized based on the AAV-EFS-NLS-CasRx-U6-DR-sgRNA_LacZ-DR sequence (Addgene Plasmid #154003 deposited by Hui Yang) ^24^. The obtained CasRx sequence was used as a template for PCR mutagenesis to introduce N2V8 mutations (A134V, A140V, A141V, and A143V)^23^. To generate donor vectors for KI into the *AAVS1* locus, the KOX1 KRAB-dCas9 gene in pAAVS1-NDi-CRISPRi (Gen1) (Addgene plasmid #73497: a gift from Bruce Conklin)^17^ was replaced with either ZIM3 KRAB-dCas9 or CasRx N2V8 using NEBuilder HiFi DNA Assembly technology (New England Biolabs), resulting in pAAVS1-NDi-ZIM3-dCas9 and pAAVS1-NDi-CasRx, respectively. The plasmids contained Cas and mCherry genes, which were linked by a P2A peptide and regulated by a Dox-inducible promoter. In contrast, the neomycin resistance genes are regulated by the endogenous *AAVS1* promoter in the event of a successful KI.

Furthermore, the PB-U6-CNCB and PB-hU6-CasRxDR1-N-CNCB vectors were used for sgRNA expression of CRISPRi and CasRx, respectively. This plasmid contained a U6 promoter-driven sgRNA expression unit, along with CAG promoter-driven Clover and Blasticidin S resistance genes flanked by piggyBac inverted terminal repeats. The sgRNA sequences are listed in the key resources table.

### Generation of human PSC lines carrying Dox-inducible Cas expression alleles

pAAVS1-NDi-ZIM3-dCas9 or pAAVS1-NDi-CasRx, in conjunction with espCas9 1.1-sgAAVS1-T2, were transfected using a P3 Primary Cell 4D-Nucleofector X Kit S (Lonza) and Program CA-137 on a 4D Nucleofector device (Lonza). Three days post-transfection, the transfectants were selected with 200 μg/mL of G418 (Wako Chemicals) until the non-transfected cells were completely eliminated. Subsequently, 1 μg/mL of Dox was added to the medium and the G418-resistant cells were cultured for 24 h. Thereafter, mCherry-positive cells were seeded at a density of one cell per well in a 96-well plate using a FACS Symphony. After 8 d of culture, colonies derived from single cells that uniformly expressed mCherry were passaged and expanded.

### Conventional PCR for genotyping

Genomic DNA was isolated from the clones using the DNeasy Blood & Tissue Kit (QIAGEN) according to the manufacturer’s instructions. Subsequently, genotyping PCR was performed using KOD One PCR Master Mix (TOYOBO) with 30 ng of genomic DNA as the template. The AAVS1-SCR-F1 and Neo-AS49 primer sets were used to confirm successful KI. Additionally, clones with KI genes in both alleles were selected using a primer set (AAVS1-WT-F2/AAVS1-WT-R2) that detected the non-KI allele. The amplicons were separated using agarose gel electrophoresis, and the images were obtained using a Gel Doc XR+ Gel Documentation System (Bio-Rad).

### Copy number assay using Droplet Digital PCR

Fifty nanograms of purified genomic DNA was digested with 2.5 U of HaeIII (New England Biolabs) in 1× CutSmart buffer, in a final reaction volume of 20 μL. Digestion was performed at 37°C for 1 h, followed by enzyme inactivation at 65°C for 20 min. The custom 20× mix of forward and reverse primers (9 μM each), along with a FAM-labeled probe targeting the neomycin (Neo)-resistance gene (5 μM), was developed by Bio-Rad. Droplet digital PCR reactions were assembled in a total volume of 25 μL containing 2 μL of the digested genomic DNA, 12.5 μL of 2× ddPCR Supermix for Probes (Bio-Rad), 1.25 μL of the FAM-labeled Neo primer/probe mixture, 1.25 μL of 20× HEX-labeled RPP30 reference primer/probe premix (Bio-Rad), and 8 μL of PCR-grade water. Droplets were generated using a QX100 Droplet Generator (Bio-Rad) according to the manufacturer’s protocol. Thermal cycling was performed under the following conditions: initial denaturation at 95°C for 10 min; 39 cycles of denaturation at 94°C for 30 s and annealing/extension at 58°C for 1 min; followed by a final enzyme deactivation step at 98°C for 10 min. Following amplification, the droplets were read using a QX100 Droplet Reader (Bio-Rad) with a CNV2 analysis setting corresponding to a two-copy reference. Absolute copy numbers were calculated using QuantaSoft software (Bio-Rad).

### Transposon-mediated gene transfer

One microgram of the sgRNA plasmid, along with 0.5 μg of a plasmid encoding a hyperactive PB transposase (hyPBase), was transfected into 5 × 10^5^ PSCs carrying Dox-inducible Cas gene alleles using the P3 Primary Cell 4D-Nucleofector X Kit S and Program CA-137 on the 4D Nucleofector device. Two days post-transfection, the transfectants were selected with 10 μg/mL of Blasticidin S (Wako Chemicals) until the non-transfected cells were completely eliminated. Subsequently, single cell-derived colonies that uniformly expressed Clover green fluorescent protein were isolated and proliferated.

### siRNA transfection

To knockdown *ATF5* and *NUP62*, a 30-pmol siRNA cocktail—consisting of three different siRNAs—was introduced into 1 × 10^5^ PSCs using Lipofectamine RNAi MAX transfection reagent (Thermo Fisher Scientific), according to the manufacturer’s instructions. Additionally, a Silencer Select Negative Control was used as a negative control. The medium was changed the following day, and 7-AAD staining and observations were performed 3 d post-transfection.

### RNA extraction and quantitative reverse transcription PCR

The cells were rinsed once with D-PBS and subsequently lysed using QIAzol reagent (QIAGEN). Total RNA was purified using a Direct-zol RNA Miniprep kit (Zymo Research) according to the manufacturer’s protocol, including in-column digestion of genomic DNA. Subsequently, complementary DNA synthesis was performed using 1 µg of total RNA with ReverTra Ace qPCR RT Master Mix (TOYOBO). Quantitative PCR analysis was subsequently performed using gene-specific primer sets (listed in key resource table) with either THUNDERBIRD Next SYBR qPCR Mix (TOYOBO) or TaqMan probe-based assays (Thermo Fisher Scientific) in combination with TaqMan Universal Master Mix II without UNG (Thermo Fisher Scientific). Amplification and signal detection were performed using a QuantStudio 5 Real-Time PCR System (Applied Biosystems). Cycle threshold (Ct) values were normalized to *GAPDH* as an internal reference gene using the comparative Ct (ΔΔCt) approach, and relative transcript levels were expressed as fold changes compared to the control samples.

### Protein analysis based on molecular size

The cells were briefly rinsed with D-PBS and lysed in RIPA buffer (Sigma-Aldrich) supplemented with a protease-inhibitor cocktail (Sigma-Aldrich). Cell lysates were clarified via centrifugation at 15,300 *× g* for 15 min at 4°C, and the resulting supernatants were collected in fresh tubes. Subsequently, the protein concentrations were determined using the Pierce BCA Protein Assay Kit (Thermo Fisher Scientific) and quantified using an EnVision 2104 plate reader (PerkinElmer), as described previously. For size-resolved and quantitative protein detection, samples were analyzed using automated capillary electrophoresis systems (Wes or Jess; ProteinSimple) equipped with 12-230 kDa separation modules (ProteinSimple). For each analysis, 2 μg of total protein lysate was loaded along with the following primary antibodies (listed in key resource table): mouse monoclonal anti-OCT3/4 (1:500; Santa Cruz Biotechnology), goat polyclonal anti-SOX2 (1:40; R&D Systems), goat polyclonal anti-NANOG (1:40; R&D Systems), goat polyclonal anti-LIN28A (1:100; R&D Systems), rabbit polyclonal anti-mCherry (1:100; Proteintech), rabbit polyclonal anti-HA tag (1:100; Proteintech), and rabbit monoclonal anti-VINCULIN (1:250; Cell Signaling Technology). Signal detection, visualization, and quantitative analyses were performed using Compass software (version 6.0; ProteinSimple).

### Immunocytochemical analysis

The cells were briefly rinsed with D-PBS and fixed with 4% paraformaldehyde (Nacalai Tesque) for 15 min at room temperature. Thereafter, the cells were incubated in a blocking/permeabilization solution consisting of D-PBS supplemented with 1% BSA, 2% normal donkey serum (Sigma-Aldrich), and 0.2% Triton X-100 (Teknova) for 45 min at room temperature. The samples were subsequently exposed to mouse monoclonal anti-OCT3/4 antibody diluted 1:200 in D-PBS containing 1% BSA and incubated overnight at 4°C. Following primary antibody incubation, the cells were washed with D-PBS and incubated for 45 min at room temperature with 1% BSA containing Alexa Fluor 647 Plus anti-mouse IgG (1:500, Thermo Fisher Scientific) and Hoechst 33342 (1 μg/mL; DOJINDO). After a final rinse with D-PBS, fluorescent signals were acquired using the BZ-X1000 fluorescence microscope (KEYENCE), and composite images were generated using BZ-X Analyzer software (KEYENCE).

### Dead cell staining

To analyze dead cells, 1.5 mL of fresh medium containing 10 µg/mL 7-AAD (Thermo Fisher Scientific) was added to the wells with cultured cells and the plate was incubated at 37°C for 10 min. Subsequently, the 7-AAD signal was visualized using the Cy5 filter of a BZ-X1000 microscope.

### RNA sequencing

Cells were lysed using QIAzol reagent, and total RNA was purified according to the protocol described eariler. RNA quality was assessed using an Agilent RNA6000 Pico Kit on a Bioanalyzer 2100 (Agilent). The library preparation and subsequent analysis were carried out following methods outlined in previous studies^42,43^. Briefly, library was prepared with Illumina Stranded Total RNA Prep Ligation with Ribo-Zero Plus kit (Illumina) using 100 ng of DNase-treated total RNA. The qualities of the libraries were evaluated using an Agilent High-Sensitivity DNA Kit (Agilent) followed by sequencing using NextSeq 2000 P2 Reagents (100 cycles) v3 (Illumina). The cutadapt-1.12 as used to trim the adapter sequences^44^. SAM tools (version 1.10)^45^ and Bowtie 2 (version 2.2.5)^46^ was used to map the reads to ribosomal RNA (rRNA) and excluded from further analysis. STAR Aligner (version 2.7.11b) was then used to align remaining reads to the hg38 human genome^47^. RSeQC (version 4.0.0) was then used to conduct quality checks on the alignment^48^. HTSeq (version 2.0.9)^49^ was used to count the reads using the GENCODE annotation file (version 35)^50^. DESeq2 (version 1.50.2) in R (version 4.5.2) was then used to normalize the obtained read counts^51^. The DESeq2 package was also used to perform Wald tests.

